# Trans-regulatory loci shape natural variation of gene expression plasticity in Arabidopsis

**DOI:** 10.1101/2024.12.20.629817

**Authors:** Mariele Lensink, Grey Monroe, Dan Kliebenstein

**Affiliations:** Department of Plant Sciences, University of California Davis, Davis CA 95616; Integrative Genetics and Genomics Graduate Group, University of California Davis, Davis CA 95616

## Abstract

Organisms regulate gene expression in response to environmental cues, a process known as plasticity, to adjust to changing environments. Research into natural variation and the evolution of plasticity frequently studies *cis*-regulatory elements with theory suggesting they are more important evolutionarily than *trans*-regulatory elements. Genome-wide association studies have supported this idea, observing a predominance of *cis*-loci affecting plasticity. However, studies in structured populations provide a contrasting image, raising questions about the genetic architecture of natural variation in plasticity. To circumvent potential statistical difficulties present in genome-wide association studies, we mapped loci underlying transcriptomic plasticity in response to salicylic acid using recombinant inbred lines generated from two random *Arabidopsis thaliana* accessions. We detected extensive transgressive segregation in the salicylic acid response, suggesting that plasticity to salicylate in Arabidopsis is polygenic. Most loci (>75%) underlying this variation act in *trans*, especially for loci influencing plasticity. *Trans*-acting loci were enriched in genome hotspots, with predominantly small effect sizes distributed across many genes. This could potentially explain their under-discovery in genome-wide association studies. This work reveals a potentially important role for *trans*-acting loci in plastic expression responses, with implications for understanding plant adaptation to different environments.

## Introduction

Plasticity can be defined as the capacity for an individual genotype to change a trait in response to external stimuli^1^. Plastic responses allow organisms to optimize their phenotypes to respond and persist in their environment. As such, plasticity can be evolutionarily advantageous by enabling a species to persevere at a new phenotypic optimum until genetic variation arises that generates a more adaptive phenotype. This model largely depends on the strength of selection and the time scale of evolutionary change ^2,3^. Furthermore, genetic variation in plasticity has been shown to be advantageous in novel environments and increase the likelihood of evolutionary rescue ^4^. Understanding how genetic variation in plasticity is genomically distributed will provide valuable insight into the regulation of response to environmental stimuli, and inform strategies to increase plant resilience to changing climates.

While plasticity affects a wide array of physiological and morphological traits, gene expression is a key tool for studying how plasticity evolves and varies. Gene expression is inherently dynamic, considered to be more directly linked to genetic changes than morphological phenotypes, and relatively easy to quantify en masse with transcriptomics ^5,6^. Thus, studies on a gene’s expression plasticity can yield more detailed mechanistic insight into how plasticity variation occurs than may be possible with a physiological or morphological trait. These benefits have led many studies to use transcriptomics to measure and map plasticity in naturally variable populations. The identified genomic loci that explain variation in transcription are called expression quantitative trait loci, or eQTL. In the context of a single gene, plasticity can be *cis*-regulated by genomic elements adjacent to or within the gene of interest, or *trans*-regulated by loci unlinked to the gene of interest. Natural variation in plasticity is theorized to be predominantly controlled by *cis*-regulators, as *cis* effects typically have less pleiotropic effects, are predicted to experience relaxed or balancing selection (as opposed to purifying selection for *trans*-regulators), and are more influential in gene expression divergence ^7–10^. In contrast, *trans*-regulatory mutations tend to be more pleiotropic with potentially deleterious effects. The resulting theory that *cis*-acting loci are under weaker purifying selection than *trans*-acting eQTL is supported by genome-wide association (GWA) studies of natural populations that identify *cis*-eQTLs as the primary identifiable basis for plasticity variation ^11–13^. Consequently, research regarding the natural variation in plasticity and its evolution often focuses on *cis*-acting changes ^14–19^.

In contrast to GWA populations, many studies using structured populations (e.g. recombinant inbred lines originating from a bi-parental cross) find mainly *trans*-acting regulation of gene expression and plasticity. In recombinant inbred strains of *Caenorhabditis elegans* and *Zea mays*, a significantly larger proportion of *trans*-acting genes had plasticity eQTL in contrast to *cis*-acting genes ^20,21^. In inbred mice, expression plasticity was also largely attributed to *trans*-acting loci ^22^. Supporting these observations was work on a small *Arabidopsis thaliana (Arabidopsis hereafter)* collection of accessions that showed the majority of plasticity variation links to coexpressed gene networks, suggesting mainly *trans*-eQTL^23^. The difference between GWA and bi-parental populations establishes a discord regarding whether variation in gene expression plasticity is predominantly controlled by *cis* or *trans* factors.

One possible explanation for the difference in the relative proportions of *trans* and *cis*-acting eQTL between structured and GWA populations is ascertainment bias, caused by the nature of *cis* vs *trans* gene regulation. *Cis* eQTL frequently have larger effects on a transcript’s expression compared to *trans* loci. GWA studies, leveraging the diversity of alleles in natural populations, are generally powered to detect loci with moderate to large-effects but are less effective at detecting loci with smaller effects. This inherent bias in GWA studies favors associations with moderate-frequency alleles that exhibit large effects ^24,25^, and detected rare alleles are likely to have an overestimated effect size ^26^. In selfing species such as *Arabidopsis*, population structure and the trait of interest are more likely to be confounded, complicating the detection of true associations in GWAS ^27^. In contrast, structured populations, by having fewer genotypes per locus and balanced allele frequencies, are better suited to find these smaller effect loci ^28–31^. The advantages and disadvantages of GWAS and QTL studies are comprehensively discussed in ^32^.

Previous work in Arabidopsis has followed this trend; GWA of transcript expression in specific conditions and their plastic response has largely found and focused on *cis*-regulation ^15,16,33–35^. To map the architecture of plasticity and assess the relative contributions of *cis* and *trans* eQTL within Arabidopsis, we used the structured Bayreuth (Bay-0) x Shahdara (Sha) Arabidopsis recombinant inbred line (RIL) population. This RIL population was treated with the defense signaling hormone, salicylic acid, or control treatment, and the transcriptome was measured at 28 hours post-treatment. Salicylic acid was chosen for plasticity analysis as it is a key hormone for controlling plant defense and exogenous application elicits responses that mirror endogenous responses. This provides a defined treatment that can induce plasticity responses similar to those in plant-pathogen interactions while minimizing physiological shifts. The Bay-0 X Sha population was chosen as these are diverse representatives of Arabidopsis with species typical salicylic acid responses ^23,36,37^. The 28-hour time point was chosen to maximize the plasticity response of gene expression while minimizing any physiological change ^23^. Using this transcriptome data, we mapped plasticity eQTL controlling variation in the SA response and show that they predominantly function in *trans* and with small effects.

## Materials and Methods

### Experimental design and data collection

We acquired transcript abundance data for the Bay-0 X Sha parental and F8 RIL lines ^38^ from a previous microarray experiment ^36^. In this experiment, the parents and 211 RILS were grown in a randomized design and treated with either a control silwet L77 (a surfactant), or 0.3mM salicylic acid (SA) in silwet L77 ^36^. The control and SA-treated plants were grown together and tissue was collected 28 hours after treatment ^23^. Growth and treatment protocols for plants are fully described in ^36^. The entire experiment was independently replicated twice. The transcriptomic analysis of the SA treatment has not been previously reported or compared to the control treatment to investigate plasticity.

The microarray used was the ATH1 Genome Array where each of 24,000 transcripts is measured using 11 probes per transcript, with each probe having 25 bases (ThermoFisher Scientific). The 11 independent and non-overlapping probes per gene are predominantly in the 3’UTR of the gene and designed using the Arabidopsis genome sequence to maximize the ability to specifically measure an individual transcript compared to any closely related sequences. Any potential cross-hybridizing probe sets are identified in the original design. This minimizes potential cross-signal caused by related sequences for other genes ^39^.

### Transcriptome Analysis

Using the expression estimate for each transcript across all the genotypes, we tested the distribution of variance linked to experimental replicates, treatment, genotype, and their interaction in both the parental accessions and RILs. This was done using a linear mixed model within the R package”R/car ^40^, with experimental replicate as a random effect while genotype and treatment were modeled as fixed effects. The sum of squares for each factor for each transcript was then calculated using a type III Anova. The average expression for each transcript within each treatment was estimated by calculating the mean value of every transcript for every individual across biological replicates. The mean values were then used to visualize how the transcriptome varies across recombinant inbred and parental lines via a principal component analysis using R/stats. This was repeated after removing transcripts that contained *cis*-eQTLs to test if the pattern was altered.

### eQTL Analysis

The average value per transcript per treatment per genotype was then used for eQTL mapping. The RILs were previously genotyped using a combination of genetic markers and the map obtained from ^37,38^. Using this map, we conducted eQTL analyses with 210 of the 211 RILS due to missing genotype information for RIL 417. For this analysis, we used the R/QTL2 package using Haley-Knott regression ^41^. eQTL analysis was performed on transcript values obtained using either silwet (control) or salicylic acid samples and as a mathematically derived value called “delta”. Delta represents an estimate of the plasticity phenotype obtained by calculating the difference in each transcript abundance across the two treatments divided by the average transcript abundance across the two treatments: (SA-SW)/[(SA+SW)/2]. This was calculated independently for each genotype for each transcript. These 3 eQTL analyses, being SA, SW, and delta, will hereafter be referred to as “expression phenotypes”. QTL locations were estimated for each transcript for each expression phenotype within R/QTL2 using a significance threshold of 2 LOD. This threshold was previously shown by permutations to be a suitable balance of type I and type II error rates ^36,37^.

The distribution of eQTLs across the genome was plotted using a sliding window with a 10 cM range and a step size of 1. For the sake of downstream analysis, transcript names were converted to the standard Arabidopsis Genome Initiative (AGI) naming system. Changes in genome predictions and gene annotations led to the removal of 1,201 genes that were measured by the microarray but are not present in the current genome annotation model. This left 88,729 eQTLs. To identify potential eQTL hotspots, permutation analysis was used to estimate a global significance threshold for QTL position within each expression phenotype (SW or control, SA, and Delta). The 95th percentile value was used for the threshold to indicate significantly enriched eQTL hotspot regions.

### Determining *Cis* vs *Trans* eQTL position and treatment conditionality

To determine *cis*-acting versus *trans*-acting QTL, the position of every gene in the *Arabidopsis thaliana* genome was interpolated into the genetic map using the Single Feature Polymorphism (SFP) marker positions from ^36^. The threshold for calling an eQTL *cis* was if the eQTL for a transcript was located 10cM in either direction of the transcript’s genomic position based on previous work ^36^. To assess an eQTLs treatment effect, we compared the map positions for eQTLs for a transcript across the control, SA, and delta expression phenotypes. If there was an eQTL for a transcript in one or more of these conditions that mapped within 10cM it was considered to be the same QTL. The eQTL was then collapsed and the average position recorded as the shared map position for further analysis. Previous work has shown that the presence of SNPs within the probes for the different transcripts on the microarray had minimal impact on the identification of eQTLs ^23,36,37,42^.

### Calculating eQTL effect size

The effect size for each eQTL for each transcript was estimated using linear regression analysis for which the predictor variable for each model was the genotype of a given eQTL across the RILs, and the response variable was the transcript abundance across the same RILs within a specific expression phenotype (control, SA or Delta). The R-squared value ascribed to genotype from each model was then used to represent the effect size of each eQTL for each transcript in each expression phenotype (control, SA, or Delta).

### Sequence comparison of Bay-0 and Sha to explore potential mechanistic origins of eQTLs

To explore the molecular mechanisms that might influence the presence and magnitude of eQTLs, we used the TAIR10 Arabidopsis Col-0 reference genome, along with Bay-0 and Shahdara assemblies from ^43^. We analyzed synteny-informed gene copy number variations using GENESPACE ^44^. For each Arabidopsis accession, we quantified the number of paralogs associated with transcripts exhibiting eQTLs across different expression phenotypes. An eQTL was considered to be located in a hotspot if it fell within 5 cM of the salicylate or jasmonate hotspot regions on chromosomes 2 and 5, respectively (refer to supplementary table 1). Due to differing gene nomenclature in the Bay-0 and Shahdara assemblies, we used the Columbia reference to identify potential *trans*-eQTLs caused by the presence or absence of paralogs within 10 cM of the eQTL loci. Additionally, we excluded 1,000 control eQTL transcripts from further comparative analysis after discovering they were fragments not representing whole genes in the TAIR assembly. The location of paralogs in centimorgans was then inferred using the same genetic map as previously described.

## Results

### Increased influence of genotype and plasticity on transcription in RILs

To understand the genetic and environmental factors controlling transcriptional variation, we modeled the relative contributions of genetics, treatment (plasticity/environment as the SA vs SW application), and the interaction of genetics and treatment (i.e. genotype x environment/plasticity) to variation in transcript abundance. This was done independently within the Parents and RIL genotypes across transcripts. In this analysis, variation between the control and SA treatments is an estimate of the plastic response to SA treatment. For each transcript, we performed linear modeling and used the results to calculate the proportion of variance for each factor. We found distinct differences between the Parents and RILs in the relative contributions of genotype, treatment, and genotype x treatment in influencing the expression of transcripts. First, most of the transcript variance in the RILs is associated with genotype and genotype-treatment while the parents’ transcript variance is predominantly in the treatment term (**Figure 1A and B**). The transcripts in the parents exhibit diverse expression patterns, with some being entirely genotype-dependent, others predominantly plasticity-dependent (the difference between SA and control) and some having no contribution from any of the experimental components. The parents also exhibit higher variability in the contribution of residuals to expression than the RIL population **(Figure 1A)**. In contrast to these variable contributions in the parents, the same transcripts in the RILs were all influenced by genotype and genotype by treatment at similar levels **(Figure 1A)**. The shift where some transcripts in the parents are mainly influenced by treatment to the same transcripts being mainly influenced by genotypes x treatment in the RILs suggests the potential for multiple polymorphic loci with opposing effect directions influencing SA response in the parents. Upon crossing and recombination, it is possible that the alleles at these loci shuffle to create new combinations in the RILs. If this is true, there should be transgressive segregation in SA responses (plasticity) in the RILs where the RILs have SA responses not found in the parental accessions.

**Figure 1.**
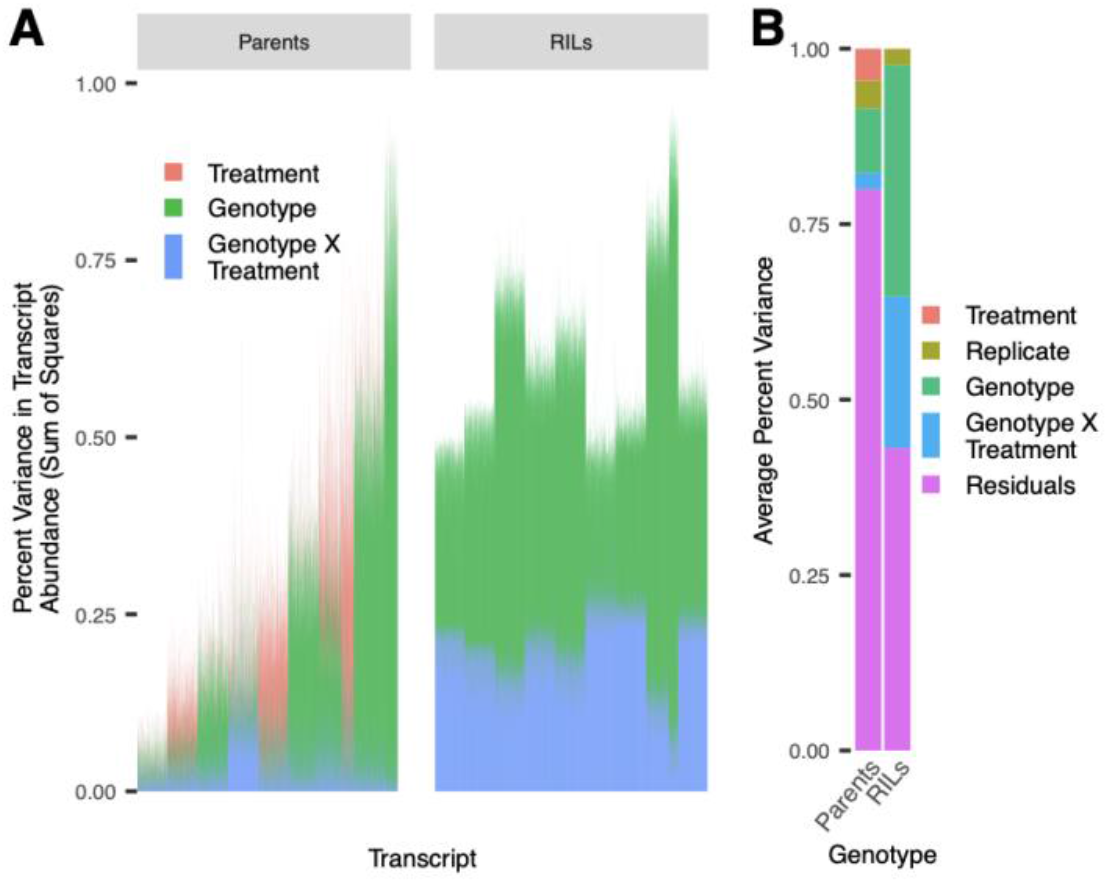
Changes in gene expression across environments are attributed to separate experimental variables between parent and RIL genotypes. **(a)** Variance in transcript abundance (sum of squares/TSS) attributable to each experimental variable for parents and RILs. Results from the type III ANOVA for each transcript were clustered into 10 variance profiles using kmeans, and 1,000 transcripts were sampled from each cluster for visualization **(b)** Average proportion of variation (sum of squares) across transcripts explained by each experimental variable.

### Greater transcriptome variation in recombinant inbred lines

In light of the differences in how genotype and plasticity influence transcripts between the parents and RILs (delta of SA response), we analyzed the overall pattern of individual transcriptomes across RILs and parents to understand general trends better. We performed principal component analysis (PCA) across all individuals first using their whole transcriptome. Results from this PCA are consistent with the idea that there is transgressive segregation for the transcriptomes within the RILs. For salicylic acid and silwet treatments, along with delta (representing a change in expression across environment, or plasticity), the RILs exhibited more transcriptome diversity than the parent lines **(Figure 2)**. This pattern implies that the generation of new genotypes also generates new expression patterns that are not present within the parental genotypes. To further visualize this, we queried individual transcripts classically associated with SA responses. Plotting each gene’s change in expression in response to treatment across individuals showed that for each of these genes, the plasticity responses of the RILs exceed the range set by the parents **(Figure 3)** which is not unexpected in segregating hybrid populations ^45^. Interestingly, this led to shifts in the sign and magnitude of SA response across RILs with genes typically induced by SA showing no change or even being repressed **(Figure 3)**. The increased variation in the global transcriptome and for expression response of known salicylic acid response genes between RILs and parental lines is evidence for transgressive segregation for gene expression plasticity within this population. Directly mapping loci and identifying opposing allelic effects between loci can validate this model.

**Figure 2.**
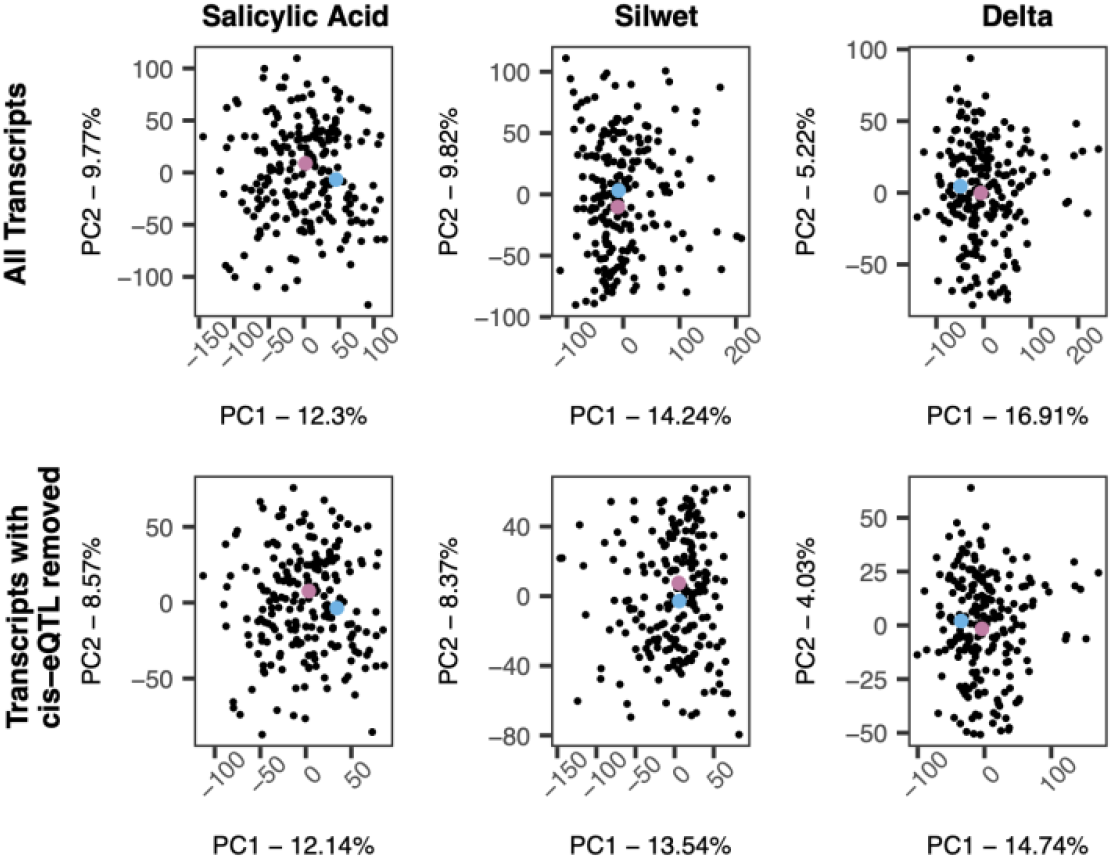
Transcriptome variation across recombinant inbred and parent lines. Principal component analysis reveals high variation in transcript abundance in recombinant inbred lines compared to parental lines in Bay-0 (blue) and Sha (pink). Analysis using the entire transcriptome is shown on the top row while transcriptome without genes continuing cis-eQTL is shown on the bottom row.

**Figure 3.**
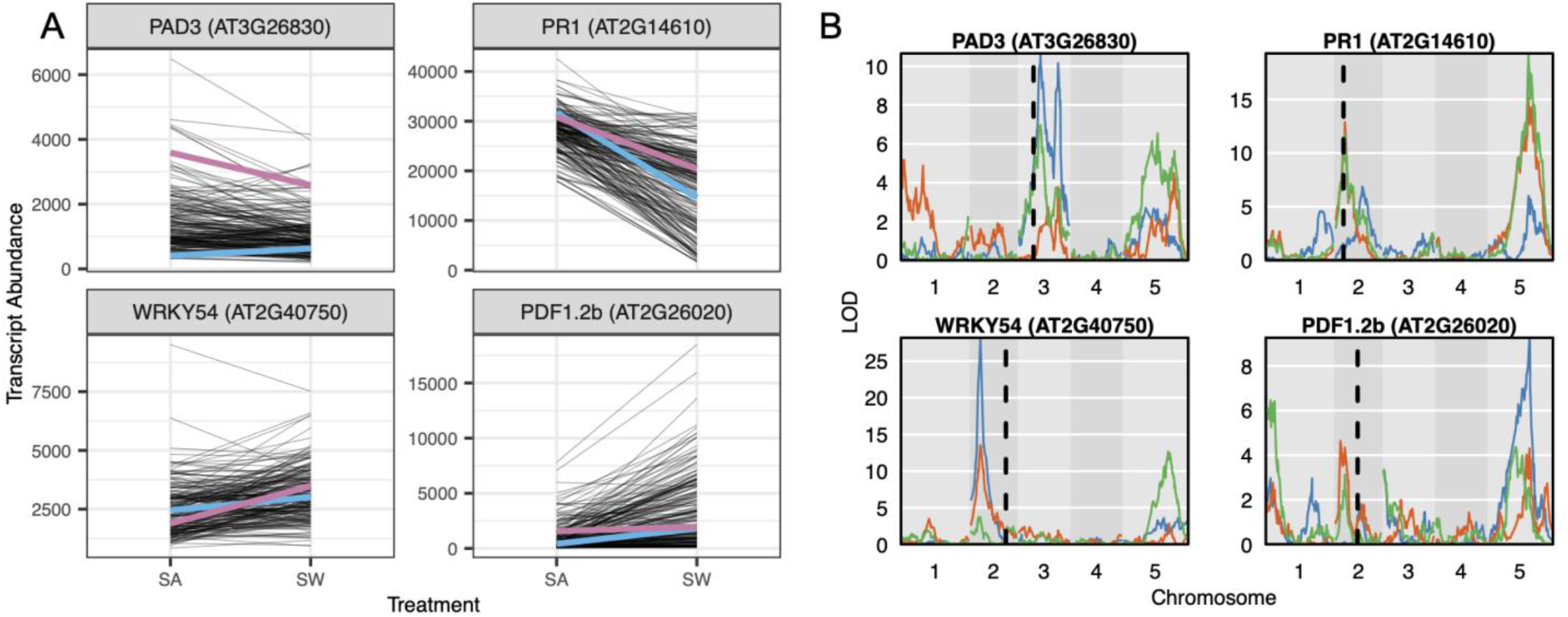
The RIL mean expression and plasticity response of classic SA response genes exceeds the range set by parent lines. **(A)** Transcript abundance of marker genes involved in salicylic acid responses across environmental treatment groups. Lines represent abundance in individual RILs (black) and the parent lines Bay-0 (blue) and Shahdara (pink). **(B)**The panels to the right show the corresponding eQTL map for each transcript.

### Trans-regulation as a major driver of gene expression plasticity

To identify the genetic loci involved in the regulation of gene expression in the context of both environment-specific expression and plasticity, we conducted three quantitative trait loci analyses on the following expression phenotypes for each transcript: expression in salicylic acid (SA), silwet (SW), and delta. We mapped eQTL for each expression phenotype and used the identified eQTL to quantify *cis*- and *trans*-regulatory eQTLS and map hotspot positions. In total, we detected 8,007 *cis* and 30,911 *trans* eQTLs for SA, 7,844 *cis* and 29,398 *trans* eQTLs for SW, and 1,278 *cis* and 11,291 *trans* eQTLs for delta. There was an average of 1.54 *trans* eQTL per transcript and 0.70 *cis* eQTL per transcript across all three expression phenotypes. For all expression phenotypes, most eQTL were *trans*-acting (∼80%) with a slightly higher fraction in the delta phenotype (90%), indicating that regulation of plasticity is highly polygenic **(Figure 4a)**.

**Figure 4.**
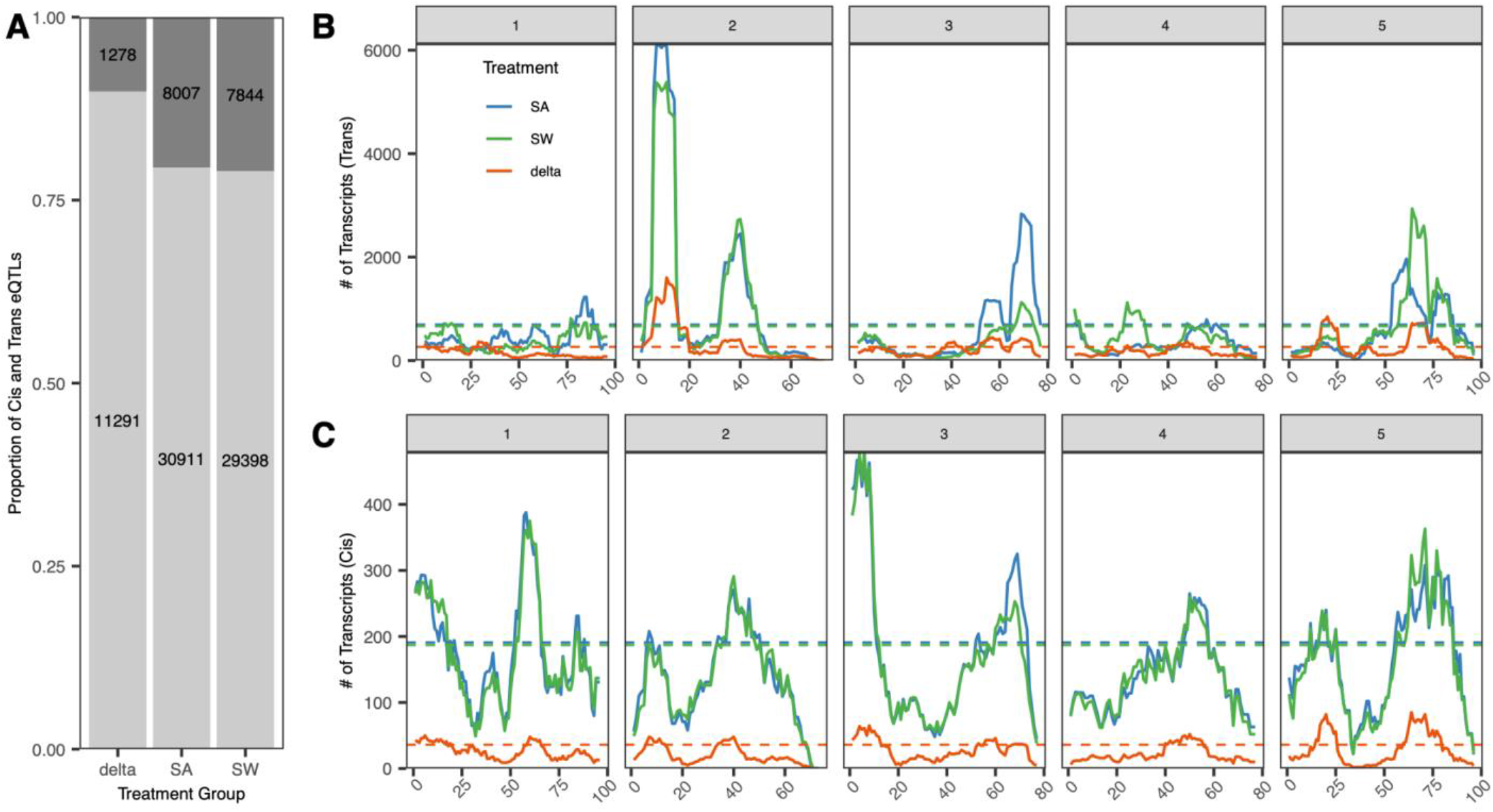
Quantification and position of cis and trans eQTLs across treatment groups. **(A)** The proportion of detected eQTL mapping to cis (dark grey) or trans (light grey) positions for each treatment group, (b) Number of eQTLs at each position (cM) across all 5 chromosomes of the A. thaliana genome for trans **(B)** and cis **(C)** eQTL. The dotted lines represent the 95th percentile value of 1,000 randomized permutations for each treatment group. The largest hot spots for all three groups are on chromosome 2. The chromosomes end at positions 100 cM for I, 70 cM for II, 80 cM for III, 80 cM for IV, and 100 cM for V

The *trans*-eQTL for all three expression phenotypes (SA, SW, delta) display similar distributions across the genome, albeit at different magnitudes. The major hotspots are located on chromosomes 2, 3 and 5, although there are secondary hotspots existing on every chromosome **(Figure 4b)**. This general pattern confirms the previous eQTL map^36^, performed on the same SW expression data. In contrast, the distribution of *cis*-eQTLs does not display any hotspot and follows the pattern of gene density across the genome. This co-localization of eQTL hotspots across the expression phenotypes implies a shared genetic architecture between plasticity (delta) and within-treatment expression variation (SA and SW). It is clear that most QTLs are *trans*-acting, especially for plasticity, and are similarly distributed across the genome for the different expression phenotypes (**Figure 4**). The presence of multiple hot spots for *trans*-eQTL, particularly for plasticity, indicates that multiple loci influence a large number of transcripts. Supporting the observation of transgressive segregation, these *trans* hotspots often have alleles with opposing directional effects for the same transcript (**Figure 3A**). This suggests that Bay and Sha parents have variation at a set of loci controlling SA responses and that the allelic variation is balanced such that the parents have the same transcriptome responses. When the allelic variation at these loci is shuffled, the parents can have dramatically different SA responses. Given that the Bay and Sha parents were randomly chosen with respect to SA signaling, the presence of transgressive variation suggests that Arabidopsis accessions may hide underlying genetic variation in SA responses more broadly ^46^.

### Plasticity is largely controlled by *trans-*regulatory loci of small to moderate effect size

We sought to characterize the effect size and relative abundance of *cis* and *trans* eQTLs existing in and across the three expression phenotypes (SW, SA, and delta). To estimate the phenotypic effects of each eQTL, we used a linear model for every eQTL detected for each transcript in each expression phenotype. We used the proportion of variance explained (R-squared) by genotype to estimate the eQTL’s effect size. To not double count eQTLs for transcripts, we compared eQTLs for each transcript across the three expression phenotypes and grouped them as a single eQTL if 2 eQTLs regulating the same transcript existed within 5.5 cM of each other across the three expression phenotypes. This showed that eQTLs influencing plasticity are mostly low to moderate effect size, both when the eQTL is exclusive to plasticity response, and when the eQTL is shared across all expression phenotypes. Additionally, eQTLs that influenced expression plasticity (delta, SA-delta, SW-delta, and SA-SW-delta) were less frequent than eQTLs found for only the non-plasticity expression phenotypes. Finally, the plasticity-associated eQTLs had greater proportions of *trans*-to *cis*-QTL, and low effect sizes compared to eQTLs that solely affected the accumulation in SA, SW, or SA-SW **(Figure 5)**.

**Figure 5.**
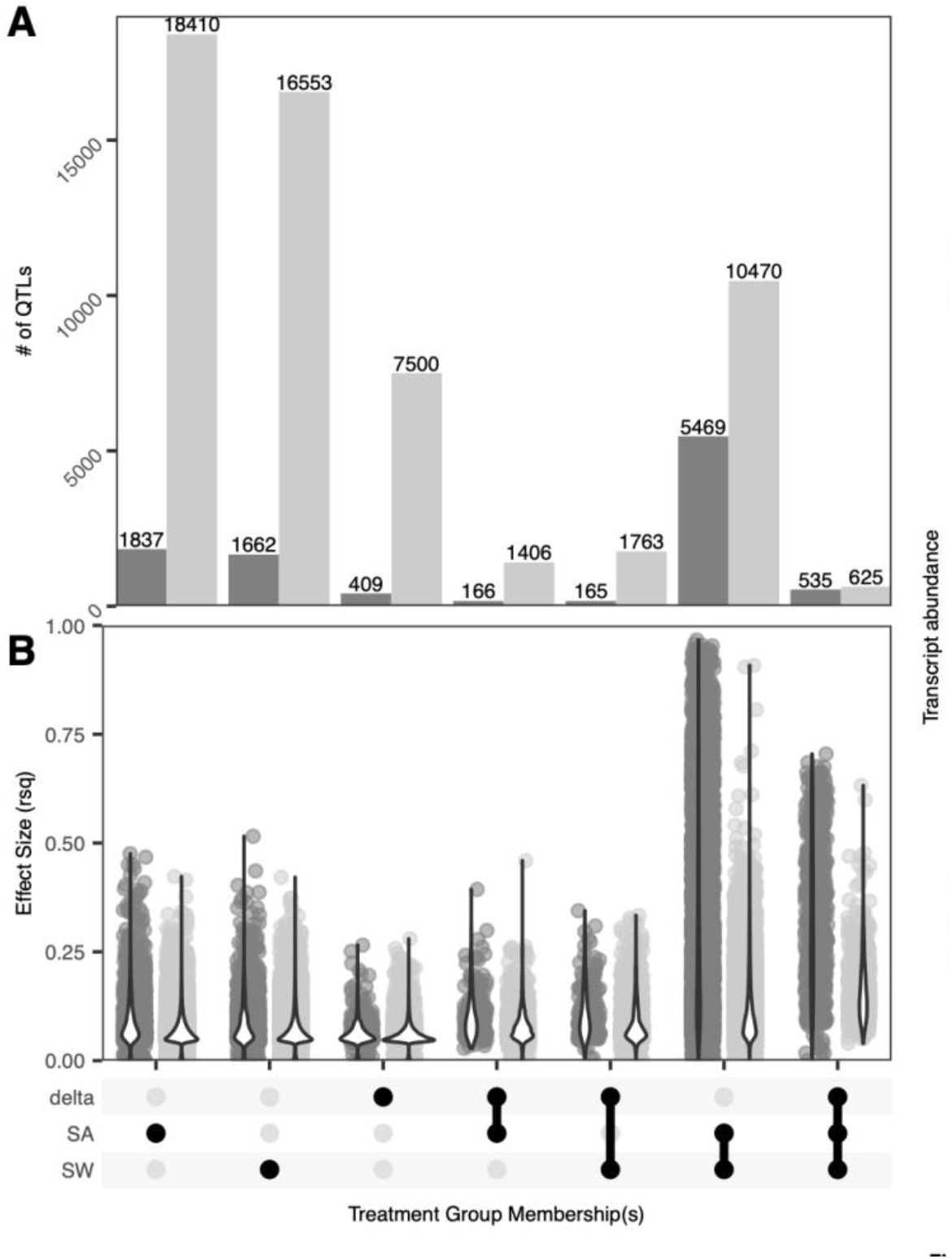
Quantity and effect size of cis and trans eQTL according to membership across treatment groups. Cis (dark grey) and trans (light grey) eQTLs present in each of the three treatments. (A) Number of eQTLs present in salicylic acid, silwet, delta, or a combination of treatment groups. (B) Effect size of eQTLs according to treatment group membership. Effect size is represented by an R-squared value. In cases where an eQTL exists in more than one group, the mean r-squared value is plotted.

It was initially puzzling that there were eQTL that linked to a transcript in SA but not SW, or *vice versa*, yet had no signal upon a plasticity/delta effect. However, plotting the effect of these eQTL on specific transcripts showed that they are likely a power-to-detect issue and likely have similar effects in both conditions and were under the significance threshold in the other condition **(Figure 6)**. In agreement with this, the eQTL effects for a given transcript showed the same magnitude and direction of change in expression across SA and SW, indicating similar effects in both SA and SW and that the original observation was caused by false negative eQTL. In contrast, transcript eQTLs linked to SA-Delta and SW-Delta effects had much stronger plasticity effects with a greater divergence in the slope of the response between the two genotypes **(Figure 6)**.

**Figure 6.**
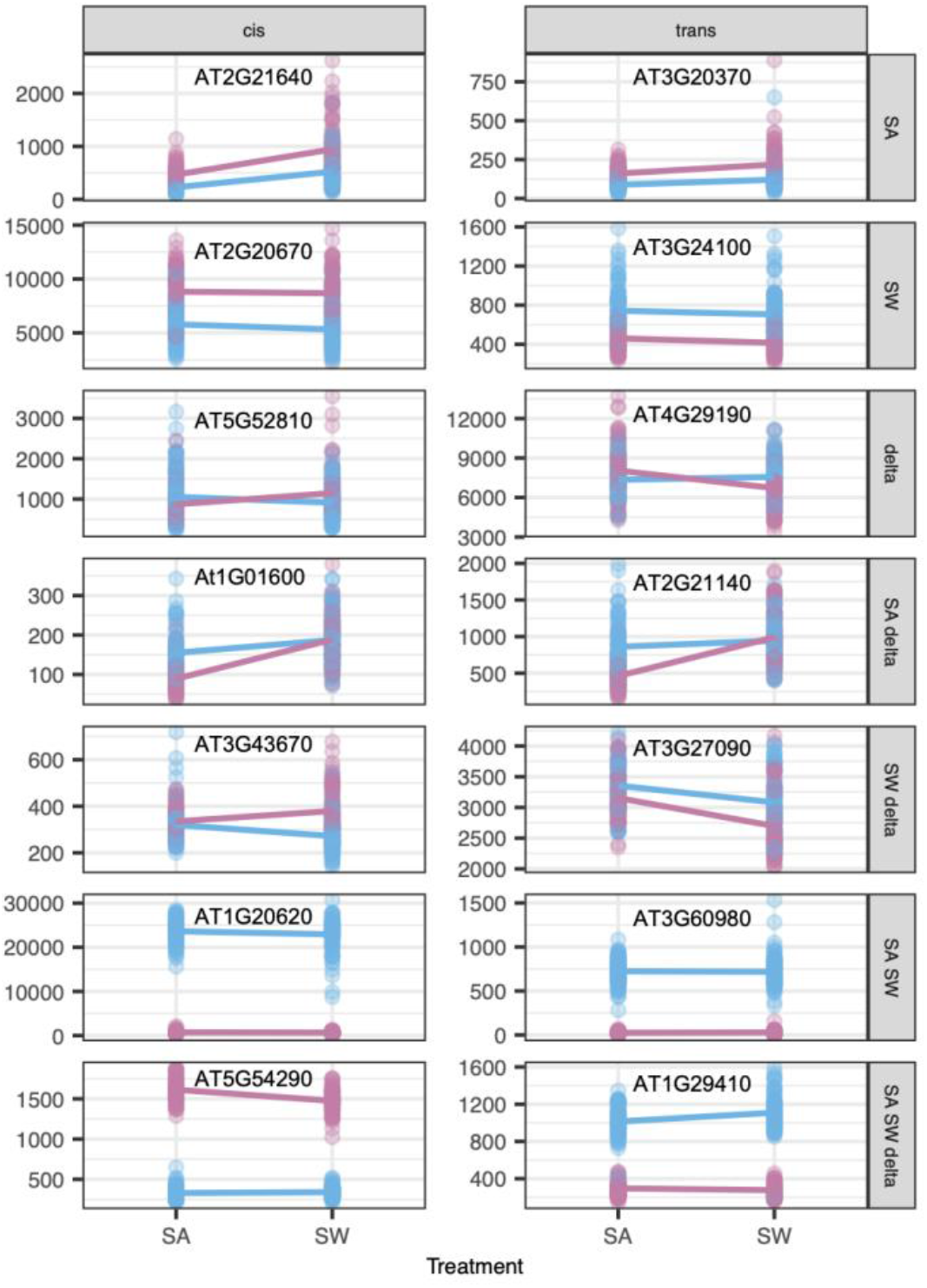
Expression across environments of high effect size eQTL according to treatment group membership. The transcript abundance for the highest effect size eQTL across type (ds or trans) and treatment group combination was plotted. Individual genotypes are colored according to the genotype (Bayreuth in blue or Shadhara in pink) at the given eQTL. Mean expression across individuals within a given genotype is represented by a solid line.

### Large-effect *trans*-eQTL

While a majority of *trans*-eQTLs are small effect size, we did identify a number of large-effect *trans* eQTL and we wanted to understand what these may represent (**Figure 7**). To better resolve these patterns and potential underlying mechanisms, we grouped all the eQTLs by their *cis*/*trans* status, presence in a hotspot and presence of a paralog at the locus. One observation from this was that eQTLs with high effect sizes often involve genes with multiple paralogs or copy number variation between accessions, especially those shared across multiple expression phenotypes. This agrees with previous observations that large effect cis-eQTL are often associated with structural variation at the gene and these show large expression differences (**Figure 5**) ^21^. The same pattern also held for large-effect *trans* eQTLs where there was a paralog to the transcript mapping to the *trans* eQTL. For example, the transcript AT1G2490 has a high-effect *trans*-eQTL across all three phenotypes, likely due to a paralog at the *trans-*eQTL that is present in Bay-0 but absent in Sha (Supplementary Table 1). Similarly, AT3G60980 has a high-effect *trans*-eQTL in SA and SW, where Bay-0 retains an extra paralog at the *trans* position, but Sha has lost the copy (Supplementary Table 1).

**Figure 7.**
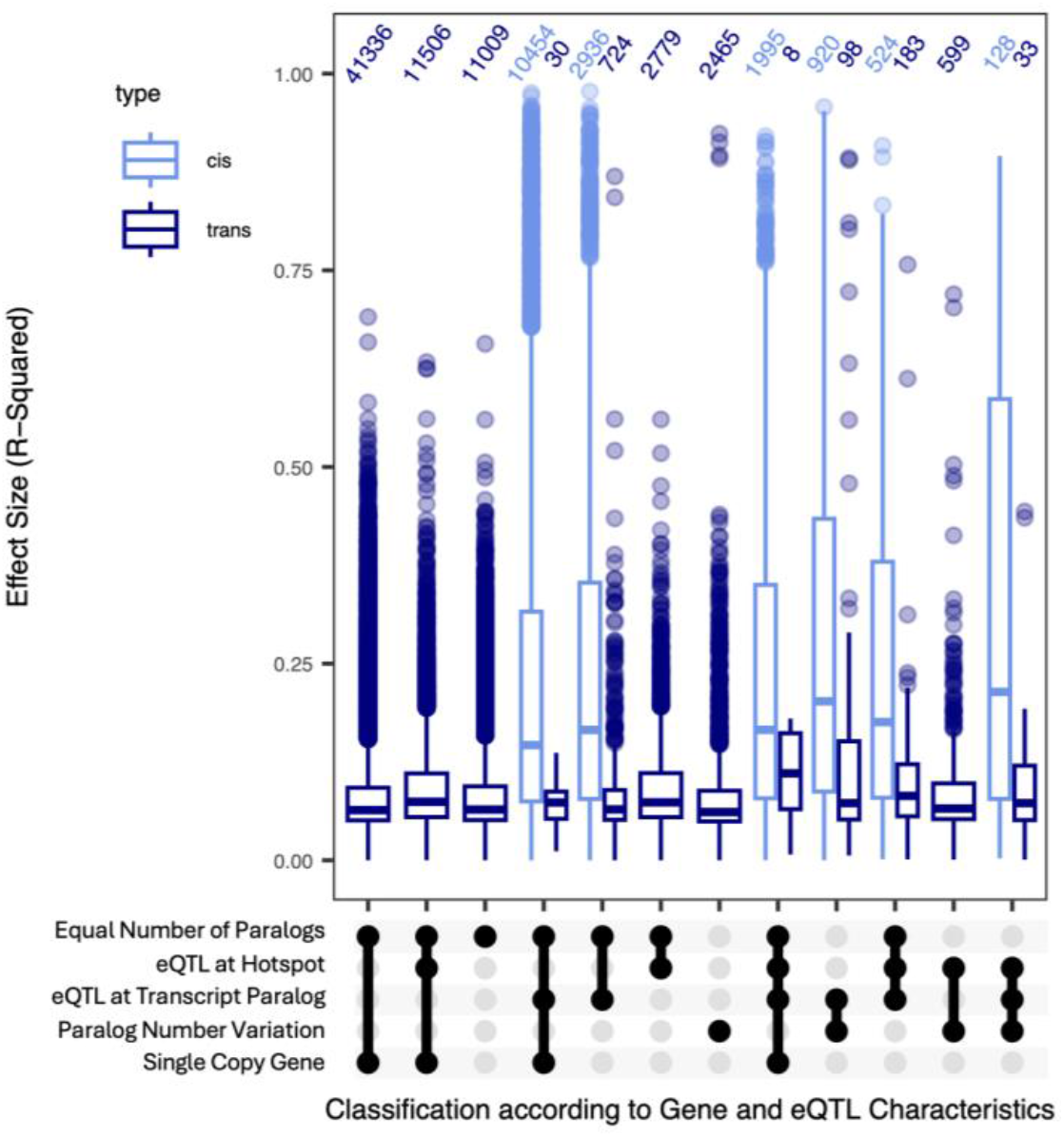
Distribution of QTL effect sizes according to genomic attributes of the transcript. Effect size of cis (light blue) and trans (dark blue) eQTLs categorized according to location of eQTL as well as paralog information of the transcript they control.

Querying this list further identified genes where *cis* structural variation in the gene family at one locus links to a *trans* effect at the other members of the gene family. There are several possible mechanisms for this. First, it may be cross recognition of the paralog by the microarray leading to a *trans* signal. Another option is that in the PAI gene family in Arabidopsis paralog variation at one locus creates a silencing mechanism decreasing expression of the other gene family members, as previously described ^47^. Additionally, some genes are known to move laterally in the genome creating an appearance of a trans signal ^37^

There were a number of large effect *trans* eQTLs that have no connection to paralogues or paralogue variation (**Figure 7**). This showed that *trans* hotspots contain large effect eQTLs affecting a number of genes. These were distributed between *trans*-eQTLs that were both within and outside of genomic hotspots. Interestingly, the presence of large effect *trans-*eQTLs within hotspots associated with plasticity suggests that pleiotropic *trans*-eQTLs are not obligated to have small effects. Assessing some of these genes showed that they included key components of the SA response and regulatory machinery (**Supplementary Figures 1 and 2, Supplementary Table 1**). Thus, while the preponderance of effects at these loci are modest to small, there can be large *trans*-eQTLs that modulate the regulatory machinery directly influencing the plasticity.

## Discussion

Transcriptomic plasticity to salicylic acid treatment within the Bay x Sha-0 recombinant inbred line population appears to be highly influenced by *trans*-regulatory eQTL. These plasticity eQTLs, because they are predominantly *trans*, tend to have a smaller effect than the *cis* eQTLs. GWA analysis are typically underpowered to find small effect loci suggesting a potential for ascertainment bias in expression plasticity studies against these plasticity *trans*-eQTL. GWA is typically powered to identify moderate to larger effect loci which in this case are *cis*-eQTL largely not associated with plasticity. This power bias could lead to an under-detection of *trans*-eQTL in GWA or other natural variation studies.

Furthermore, previous studies have demonstrated that trans-regulatory variation can influence adaptive evolution within species ^48,49^.

This raises the potential that GWA studies have significant amounts of undetected *trans*-regulatory loci influencing gene expression and suggests that incorporating structured populations into plasticity studies will be key to better resolving the regulatory landscape of plasticity variation and helping to develop better models of how plasticity variation may evolve.

Salicylic acid signaling is considered to be a highly conserved core component of plant responses to pathogen attack. This model was supported by previous studies identifying minimal variation in SA responses across Arabidopsis accessions ^46^. In agreement, we observed minimal variation in the SA transcriptome response between the Bay and Sha parents (**Figures 1 and 2**). In contrast, the RIL progeny displayed extensive genotype x SA treatment variation across nearly the entire transcriptome, and this mapped to a defined set of *trans*-eQTL hotspots **(Figures 1 and 4)**. These hotspots were pleiotropic in influencing the SA response of 100s to 1000s of transcripts **(Figure 4)**. Because these loci had opposing directional effects on the response to SA, they created a situation whereby the parental accessions had largely the same SA response while the RIL progeny showed extensive transgressive segregation in SA response (**Figure 3**). This shows that the minimal phenotypic plasticity to SA in Arabidopsis accessions hides genetic variation underlying the SA response, further supporting the need for structured populations in assessing plasticity variation within a species.

Previous research has observed that transgression is relatively common across plants and influences a diverse set of traits. The likelihood for transgressive segregation can increase for traits under constraining selection like a key defense response and also within selfing plant populations such as Arabidopsis ^45^. Population structure limitations on gene flow can also create partitions that allow for the evolution of transgressive traits. Differentiating between these models would require the development of a number of structured populations across the species to better sample the potential for transgression to be a common aspect of typically assumed conserved signaling pathways. This is probably necessary to fully understand the genetic architecture of plasticity because extensive transgressive segregation across Arabidopsis accessions would influence the interpretation of GWA ^50^. Transgressive variation and constraining selection would hide causal genetic variation that influences the trait and may be under locally adaptive selective pressures. Observations from this structured population suggest that GWA studies of transcriptomic plasticity may need careful assessment. (1) Plasticity eQTL are biased towards small effect *trans* loci which could be undetected in GWA studies. (2) Large effect plasticity *trans* eQTL can be linked to paralog variants suggesting they are actually *cis* in causation. (3) Plasticity can display transgressive segregation that can constrain GWA ability to identify causal loci. While this is from a single structured population, it was randomly chosen with regards to the traits in question and the parents represent a broad sampling of Arabidopsis genetic diversity. While it will take additional structured populations to assess this potential more broadly, it does suggest there is a need to conduct these structured population studies more broadly to better understand the results from GWA collections.

