## Supplemental Figures for "Trans-regulatory loci shape natural variation of gene expression plasticity in Arabidopsis"

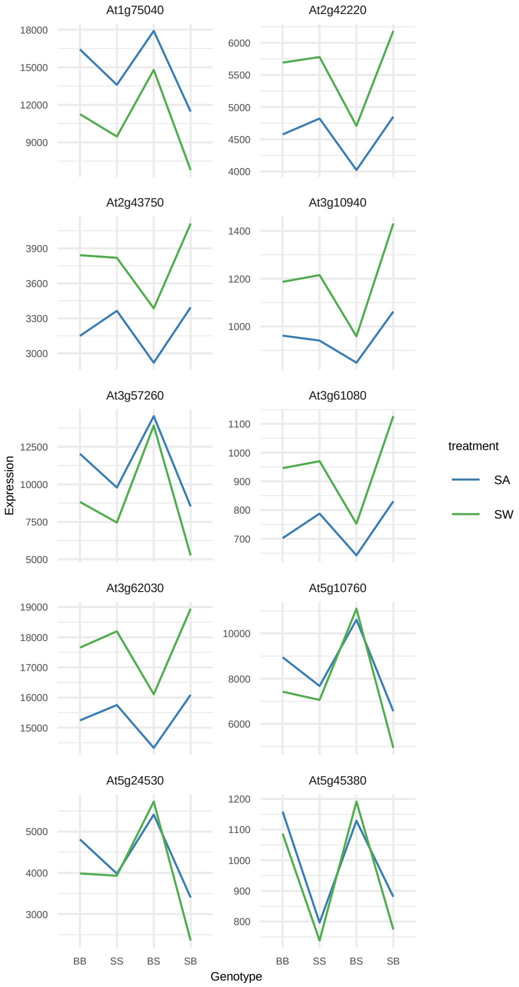


**Supplementary Figure 1. Expression of transcripts with SA hotspot eQTLs on Chromosome 2 under salicylate and silwet treatment for parental and recombinant genotypes** The x axis is the genotype of RILS at the location of the SA and JA hotspots. For example, BB represents RILs with the Bay-0 genotype at both locations, where SB is RILs with the Sha genotype at the chromosome 2 hotspot and Bay-0 at the chromosome 5 hotspot.

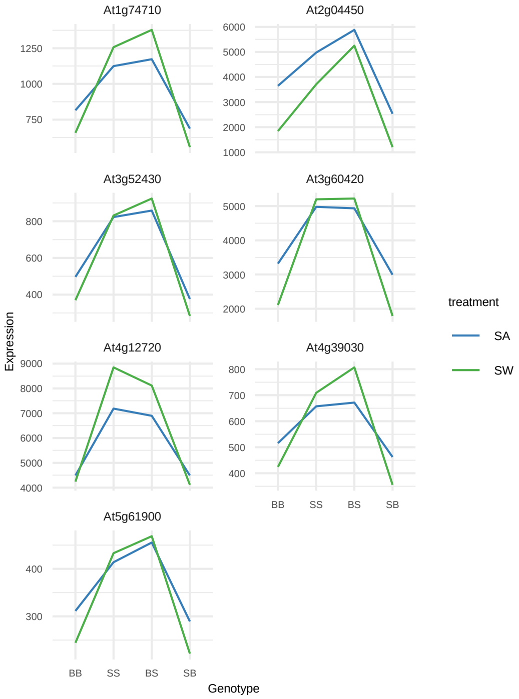


**Supplementary Figure 2. Expression of transcripts with JA hotspot eQTLs on Chromosome 5 under salicylate and silwet treatment for parental and recombinant genotypes** The x axis is the genotype of RILS at the location of the SA and JA hotspots. For example, BB represents RILs with the Bay-0 genotype at both locations, where SB is RILs with the Sha genotype at the chromosome 2 hotspot and Bay-0 at the chromosome 5 hotspot.
